# Major functional bias for mitochondrial complexes in genome-wide CRISPR screens

**DOI:** 10.1101/2020.08.31.273730

**Authors:** Mahfuzur Rahman, Maximilian Billmann, Michael Costanzo, Michael Aregger, Amy H. Y. Tong, Katherine Chan, Henry N. Ward, Kevin R. Brown, Brenda J. Andrews, Charles Boone, Jason Moffat, Chad L. Myers

## Abstract

We present FLEX (<u>F</u>unctiona<u>l</u> <u>e</u>valuation of e<u>x</u>perimental perturbations), a pipeline that leverages several functional annotation resources to establish reference standards for benchmarking human genome-wide CRISPR screen data and methods for analyzing them. We apply FLEX to analyze data from the diverse cell line screens generated by the DepMap project. We identify a dominant mitochondria-associated signal, which our time-resolved CRISPR screens and analysis suggests may reflect screen dynamics and protein stability effects rather than genetic dependencies.

---

CRISPR-based screening techniques have become a central instrument for systematic investigation of gene function. At the forefront of such efforts, the Cancer Dependency Map (DepMap) effort aims to catalogue genetic dependencies of all human genes across a range of cultured cell lines spanning various tumor entities. To date, the loss-of-function fitness effects of 17,634 genes have been measured in 563 cell lines (19Q2 data release) ^1 2^. This data provides a comprehensive and easily accessible resource for biological hypothesis generation. Several studies have developed computational methods to systematically derive functional information from this data including inferring genetic interactions ^3^ or functional relations by utilizing co-essentiality profiles ^4 5 6 7^. In more technical terms, such methods infer pairwise gene-gene relations. Despite the wealth of data and a diversity of methods for processing CRISPR screening data, we lack standard benchmarks for evaluating their ability to extract functional information, which ultimately limits our progress in establishing the best practices for analyzing CRISPR screens.

Here we developed FLEX (<u>F</u>unctiona<u>l</u> <u>e</u>valuation of e<u>x</u>perimental perturbations), a pipeline to evaluate functional screening data or algorithms designed to improve scoring or interpretation of such data. FLEX derives reference standards from diverse genome-wide functional resources such as CORUM complexes ^8^, curated pathways ^9^, GO Biological Processes (BP) ^10^, and genomic data-derived functional networks ^11^. It then uses these reference standards to (i) generate summaries of precision-recall (PR) performance on a global and local scale by assessing the degree to which genetic dependency profiles capture known functional relationships (ii) investigate underlying functional diversity driving the observed PR performance, and (iii) report a diversity-normalized PR statistic that highlights both the quality and functional diversity of functional relationships captured by a dataset of interest. FLEX is available as an R package.

We deploy FLEX to evaluate the capacity of the DepMap CRISPR knockout co-essentiality profiles to recover complex, pathway, and biological process co-membership of all genes in the human genome (Figure 1a, Figure S1). Precision-recall (PR) statistics showed that co-essentiality profiles recapitulated many known functional relationships - for example at a precision of 0.5, 3348 true positive (TP) co-complex pairs from the CORUM complex standard were identified based on pairwise Pearson correlation coefficients derived from the DepMap dataset (Figure 1b). FLEX uses PR statistics to account for the strong class imbalance typically observed in functional genomics data, where the number of positive events (true functional relationships) is much smaller than the number of negative events (unrelated pairs) ^12^. While PR statistics provide a general quantification of functional information, they do not provide insight into the diversity of functional information captured by a particular dataset. To understand how individual protein complexes contribute to overall performance, we assessed the area under the PR curve (AUPRC) for each individual complex as well as the fraction of TP pairs mapping to each complex at different levels of precision (Figure 1c, d). Strikingly, we found that only two of 1697 complexes in the CORUM standard, the electron transport chain (ETC) I holoenzyme and the 55S mitochondrial ribosome dominate the strongest correlated gene pairs, contributing ~76% of the 3348 TP pairs at a precision of 0.5 (Figure 1c). Consistent with a predominant functional signal contributed by theses complexes, exclusion of the ETC and 55S mitochondrial ribosome annotations from the 1697-complex standard, but not removal of other large complexes or small complexes with high AUPRC, vastly reduced global PR performance of the DepMap (Figure 1e), suggesting that caution needs to be taken in interpreting such global evaluations.

**Fig 1:**
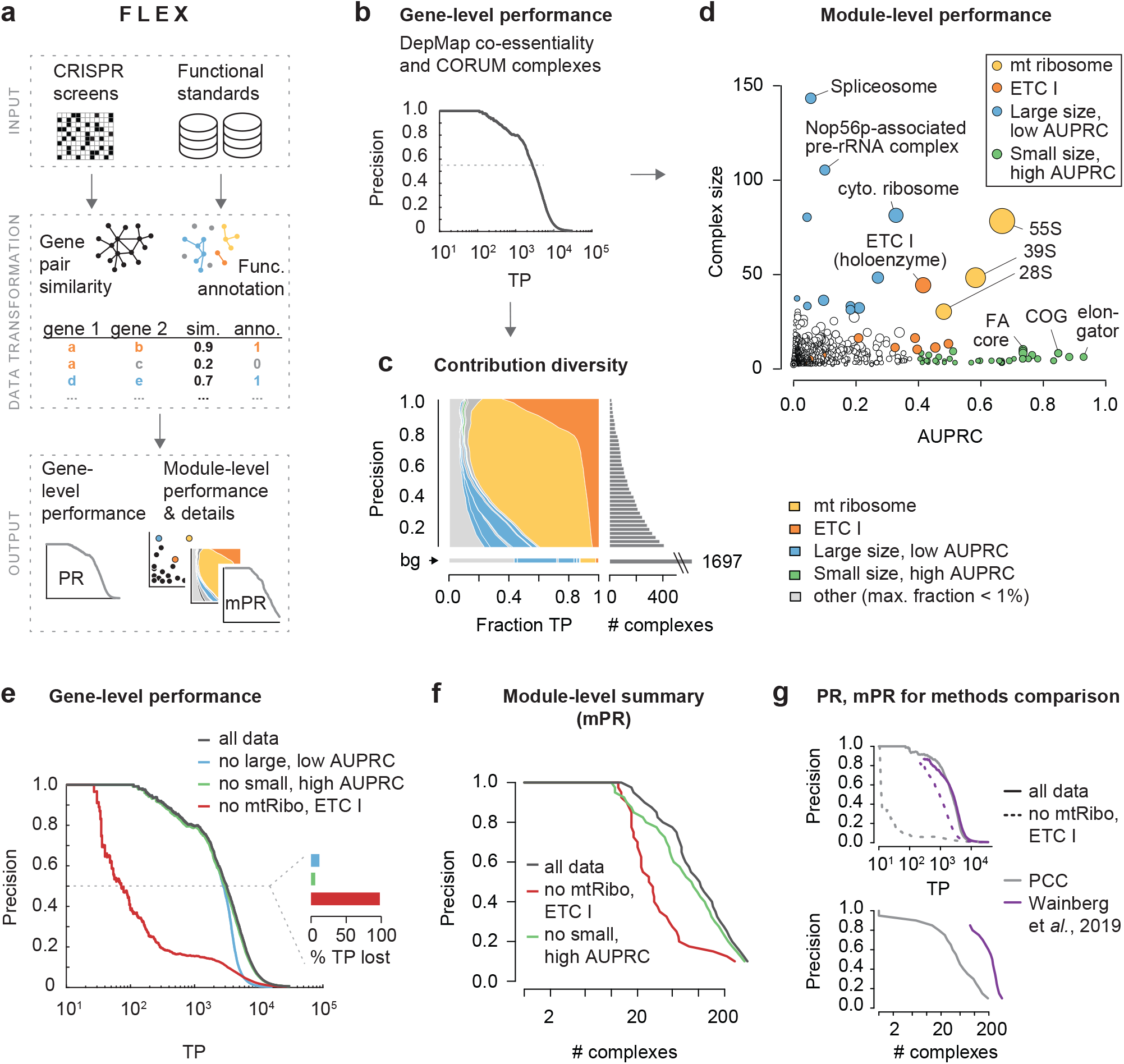
FLEX reveals mitochondrial bias in functional CRISPR/Cas9 screening data. **a**, FLEX inputs a CRISPR screening dataset and functional reference standards to compute gene-level performance and module-level (e.g. protein complex) performance summaries (see Fig. S1 for details). **b**, Precision-recall (PR) performance of gene-gene co-essentiality profile correlation using the CORUM complex standard to define true positives (TP). Pearson correlation coefficients (PCC) are computed between CERES score profiles across the 563 19Q2 DepMap screens for all possible gene pairs. **c**, Contribution diversity of PR performance using the CORUM complex standard. Shown are the fraction of TP pairs for CORUM complexes (distributions across the x-axis) at different precision cutoffs (down the y-axis). The minimum number of complexes to cover the complete set of TPs is shown (see Methods for details). Complexes with a fraction smaller than 0.01 (1%) at any precision are collectively shown in light grey. The background (bg) functional diversity represents the distribution of categories across the entire reference standard (i.e. the expected distribution in a random selection of gene pairs). The number of complexes required to cover TPs at a given precision are plotted on the right. Highlighted complexes are defined in d. **d**, Size and individual CORUM complex PR performance. Area under the PR curve (AUPRC) was computed per complex. Dot size corresponds to the mean within-complex CERES profile PCC, adjusted by the standard error. Protein complexes with at least 30 members (genes) are defined as large, otherwise small. Complexes with an AUPRC of at least 0.4 are defined as high AUPRC, otherwise low. All sub-complexes mapping to the ETC I or 55S mitochondrial ribosome are shown in the respective color. **e**, PR performance of gene-gene co-essentiality profile correlation (see b). Black line shows complete data, colored lines show the performance after sets of complexes (defined in c) were removed from the data and standard. The inset barchart shows the percentage of TP lost at a precision of 0.5 after either set of complexes is excluded. **f**, Module PR (mPR) curve summarizes performance at the module level (here, CORUM protein complexes), and thus, represents a more robust evaluation metric. The number of complexes (x-axis) captured at a minimum recall are plotted at each corresponding precision cutoff (y-axis) (see Methods for details). **g**, Comparison of two methods measuring co-essentiality in the DepMap using PR and mPR plots. The method proposed by Wainberg and colleagues is compared to the standard PCC-based method (top). The well-balanced coverage of complexes is shown after their ETC-related complex exclusion (dotted lines, top) as well as in the mPR curve (bottom).

Complexes such as the ETC and the 55S mitochondrial ribosome dominate these global evaluations because they are well-captured by profile similarity in the DepMap data, as supported by focused PR analysis of gene pairs associated with only genes in these complexes (Figure 1c-e, Figure S2a-d, Figure 3a-d), but due to their large size, they contribute a large number of pairs. To enable functional evaluations of CRISPR screen data that are less influenced by well-performing, large gene sets, we implemented in FLEX an additional, complementary metric, termed module-level Precision-Recall (mPR) performance. The mPR measure is computed by first calculating a contribution matrix, which measures the number of true positives at different precision levels for each gene set in the functional standard (e.g. each complex for the CORUM standard). These results are then summarized across all gene sets, but each gene set is allowed to contribute only a single observation, thereby stabilizing the contribution of large versus small gene sets to the evaluation (Figure 1f). We emphasize that each of these complementary FLEX visualizations is produced by default for any dataset evaluated by the pipeline; considering all of them collectively is important to gain an accurate perspective of the functional information captured by a given dataset.

FLEX enables objective benchmarking of methods for scoring or processing CRISPR screen data, several of which have been recently published specifically for the DepMap ^4 6 7^. As an example, we used FLEX to compare an earlier DepMap data release (18Q3) to a later release (19Q2), which was based on an improved CERES score ^2^, and found that the later release shows substantial improvement in capturing functional relationships and a greater functional diversity in the relationships captured (e.g. ETC-related complexes are less dominant) (Figure S2e-h). Using FLEX to evaluate a collection of published methods for analyzing DepMap co-essentiality profiles^4 6 7^, we found substantial dependence on the mitochondrial complexes in all of them, with the notable exception of the algorithm published by Wainberg and colleagues ^7^, which captures functional relationships with a much greater functional diversity than other methods (Figure S4, Figure S5). This superior performance with respect to functional diversity may result from accounting for covariance among cell lines, which is a key feature of the method by Wainberg and colleagues but not others, and this difference is clearly highlighted by FLEX's mPR metric (Figure 1g, Figure S5).

To further explore the functional signal contributed by ETC-related complexes in DepMap co-essentiality profiles, we compared dependency data from 149 cell lines in the 19Q2 DepMap that had been screened both at the Broad Institute (hereafter referred to as Broad DepMap) and the Sanger Institute (Sanger DepMap). Since we also observed a strong signal for the ETC V complex (Figure S3a), we hereafter consider ETC I, V and the 55S mt ribosome and refer to them collectively as ETC-related complexes. While the dependency profiles for the same cell lines generally agree across these two datasets ^13^, we found the ETC-related genes to be among the protein complexes exhibiting the strongest differences between them, with the Broad DepMap consistently measuring stronger drop-out phenotypes for these genes (Figure 2a). Assay length is a major difference between Broad and Sanger DepMap screening protocols (Broad screens are conducted over 21 days while the Sanger screens are completed over 14 days) ^13^, and thus, we reasoned that the difference in ETC-related genes' phenotype observed in the Broad and Sanger screens may be related to screen sampling times. Specifically, we hypothesized that the rate at which functional proteins are cleared from the cell after successful gene disruption may impact phenotypic penetrance over the course of a screening experiment. In other words, the growth phenotype associated with disruption of an essential gene would only be observed after the corresponding essential protein is mostly depleted from the cell population. Thus, in the case of a highly stable essential protein, cells may need to be cultured for a longer period of time after gene disruption to observe the resulting growth defects. Consistent with this hypothesis, we found that protein complexes with significantly more severe fitness defects in the Broad DepMap screens (FDR < 5%) tend to be more stable (z-score > 1; p = 0.001, hypergeometric test), based on analysis of available protein half-life data derived from monocytes, B cells and hepatocytes ^14^. Strikingly, the ETC I and V complexes showed the highest protein stability of any complex in the CORUM standard (Figure 2b, Figure S9a-c). In contrast, the 55S mitochondrial ribosome had a protein half-life comparable to the median complex half-life. The phenotypic delay observed for the 55S ribosome may also be related to high remaining protein levels of the ETC, which acts downstream of the 55S ribosome. More specifically, the phenotypic effect of the 55S ribosome perturbation is a result of its impact on ETC complex disruption (i.e. the ETC complex is epistatic to the 55S ribosome in this context), which may explain why it exhibits similar dynamics in the context of a CRISPR screen.

**Fig 2:**
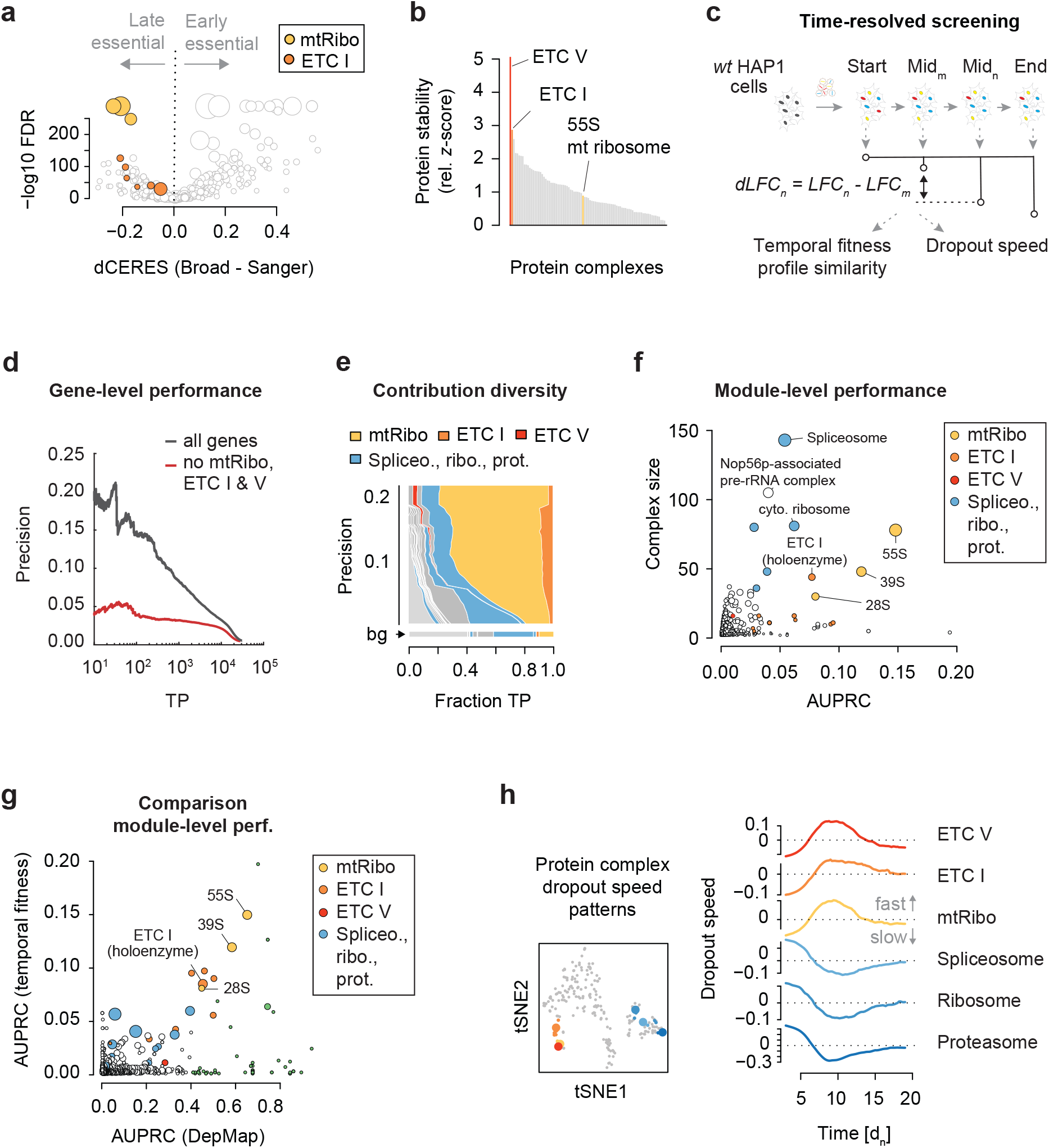
Delayed ETC CRISPR/Cas9 fitness phenotypes create within-complex co-essentiality. **a**, Protein complex-level differences in fitness effects between the Broad and Sanger DepMap screens. The 149 cell lines and 16,464 genes common to both data sets are compared. For each CORUM complex, the median differential CERES score (x-axis) and a paired Wilcoxon rank sum p-value with BH-correction are shown. Mitochondrial ribosome (yellow) and ETC I (orange) subcomplexes are highlighted. **b**, Protein stability of CORUM complexes. Protein half-life data was taken from B cells, hepatocytes, and monocytes, and summarized on the CORUM complex level. Half-life data were z-transformed, and the minimum z-score set to 0 to emphasize large z-scores. Complexes for which at least 5 members contributed data across the three cell lines are shown. **c**, Scheme of time-resolved genome-wide CRISPR/Cas9 screens in HAP1 cells. Temporal fitness profile similarity was estimated by computing the pairwise PCC between genes with 32 unique measurements across time. The dropout speed was derived from profiles interpolated from the 32 measurements after correcting for maximal dropout effects (see Methods). **d**, Precision-recall (PR) curve showing HAP1 temporal fitness profile similarity performance using CORUM complexes as a pairwise functional standard. Black line shows complete data, red line performance after ETC I, V and mitochondrial ribosome (ETC-related complexes) are removed from the data and standard. **e**, Contribution diversity of HAP1 temporal fitness profile similarity PR performance using the CORUM complex standard. Shown are the fraction of TP pairs for CORUM complexes (distributions across the x-axis) at different precision cutoffs (down the y-axis). The minimum number of complexes to cover the complete set of TPs is shown (see Methods). Complexes with a fraction smaller than 0.01 (1%) at any precision are collectively shown in light grey. The background (bg) functional diversity represents the distribution of categories across the entire reference standard (i.e. the expected distribution in a random selection of gene pairs). **f**, Module-level performance of HAP1 temporal fitness profile similarity shows CORUM complex size and AUPRC. Dot size corresponds to the mean within-complex similarity, adjusted by the standard error. All sub-complexes mapping to the ETC-related complexes are shown in the respective color. **g**, Comparison of module-level performance between Broad DepMap co-essentiality and temporal fitness. AUPRC measures the performance of each dataset in reconstructing CORUM complex co-memberships. Dot size is proportional to complex size. **h**, Dropout speed for ETC-related and other selected essential complexes. A positive dropout speed indicates faster relative dropout, while a negative dropout speed indicates slower dropout. tSNE embedding groups CORUM complexes with similar dropout speed (see Methods). The six selected complexes on the right are indicated in the tSNE plot (large colored dots) and sub-complexes are labeled with matching colors.

To test if temporal drop out patterns that are dependent on assay length and protein stability could contribute to the high similarity of ETC-related co-essentiality profiles, we performed several CRISPR/Cas9 screens in a single cell line (HAP1 cells) and measured gene essentiality at multiple different time points over the course of the screen (Figure 2c). Specifically, we sampled cells every 3 to 4 days following initial infection and compared the abundance of guide RNAs (gRNAs) targeting a particular gene at a given time point relative to the starting gRNA abundance for the corresponding gene. Applying this approach, we generated dynamic essentiality profiles, derived from seven biological replicate screens, for each of ~18,000 genes targeted by our genome-wide TKOv3 gRNA library (Figure 2c, Figure S6a-c). Similar to observations from the Broad DepMap, we found that genes with similar time-resolved essentiality profiles, derived from a single HAP1 cell line, tended to be functionally related and often annotated to the same protein complex or biological pathway (Figure 2d-f). Strikingly, using FLEX to dissect the observed functional performance revealed that the same ETC-related complexes were responsible for the majority of the functional associations derived from our dataset (Figure 2d-g, Figure S7a-d). We note that other large essential protein complexes with shorter protein half-lives (e.g. the 26S proteasome or the cytosolic ribosome) drop out relatively rapidly when targeted (within 5 days post puromycin selection) while the ETC-related complex members take substantially longer, dropping out between 6 and 13 days post puromycin selection (Figure 2h, Figure S8). This observation is consistent with phenotypes for these complexes observed in earlier CRISPR/Cas9 screens ^15^ and RNAi-based screens ^16^.

Given the delayed phenotype of protein complexes with high protein half-lives, we reasoned that the time at which a CRISPR/Cas9 screening experiment was sampled could affect the measured dependency on a particular gene target, and vice versa, that the measured dependency may reflect differences in effective sampling time. To test this, we first sorted the 563 cell lines in the Broad DepMap using the median CERES score for ETC-related complexes. As expected, while genome-wide CERES scores for each cell line exhibited comparable ranges (Figure 3a, grey), ETC-related CERES scores strongly varied across cell lines (Figure 3a, red, Figure S10a-c). Furthermore, when we added the data from the 149 Sanger DepMap screens that overlapped the Broad DepMap, a matched comparison of the ranks for those 149 showed lower ranks (weaker ETC signal) relative to the corresponding Broad screens (Figure 3b). We further tested how different time points of a single cell line, HAP1, would rank within the Broad DepMap collection of screens based on the strength of the ETC-related phenotype. We leveraged 27 time points measured in 7 independent genome-wide screens taken between 6 and 19 days post puromycin selection (Figure S6a-c) (see Methods). We found that those HAP1 screen time points spanned the range of Broad DepMap screen ranks, with early time points showing weaker dependency on ETC-related genes (Figure 3c). Notably, the strength of the ETC-related dependency itself predicted HAP1 screen timepoints with reasonable accuracy (r = 0.61, p = 0.0007). This was not true of the dependency scores for other essential complexes such as the 26S proteasome (r = −0,22, p = 0.28), the spliceosome (r = −0.01, p = 0.96) or the cytosolic ribosome (r = 0.37, p = 0.055) (Figure S8d-f). Together, this suggests that the strength of ETC-related fitness phenotypes is able to accurately recover the effective length of time a screen was cultured before gRNA abundance was quantified.

**Fig. 3:**
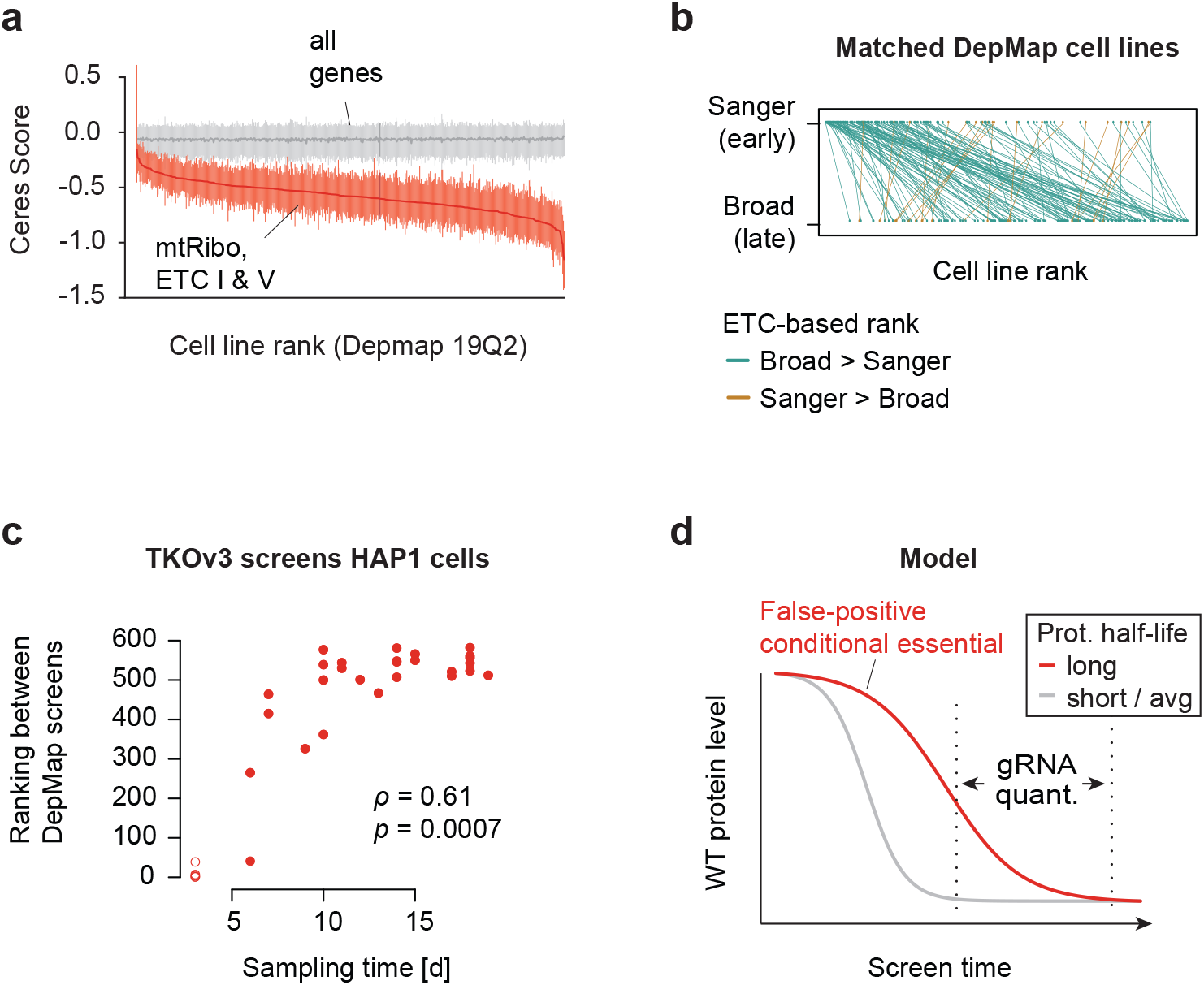
CRISPR screen sampling time and protein level change. **a**, Broad DepMap genome-wide CRISPR/Cas9 screens ranked by the median CERES score across the ETC-related complexes. The middle red line indicates the median, the vertical lines the 25% and 75% quantiles of a given screen. Grey lines represent the same metrics for all genes in the genome. **b,** Pairwise comparison of Broad and Sanger DepMap screens based on their median CERES score of ETC-related complexes. Highlighted are 149 cell lines common to both data sets. To rank those cell lines, Sanger data from those 149 screens was added to the 563 Broad DepMap screens and all screens were ranked. Green lines indicate a higher ranking of the Broad screen (assay length 21 days) and brown a higher ranking for the corresponding Sanger screen (assay length 14 days). **c**, Rank of HAP1 time course genome-wide screens in the Broad DepMap screens based on the adjusted median ETC-related LFC. HAP1 screens were performed with the TKOv3 library, and LFC values were adjusted by centering non-essential genes around 0 and core essential genes around −1 (see Methods). HAP1 screens sampled at T3 are shown as circles indicating that they have not been used for computing the Wilcoxon rank-sum correlation coefficient (see Methods for details). **d**, Wildtype protein abundance of two protein complexes is schematically displayed over the course of a CRISPR screen. The measured phenotype (e.g. gRNA abundance as a proxy for cell fitness) depends on the presence of a sufficient amount of protein to fulfill a cellular function. Stability of proteins, the rate of cell doublings that redistribute residual protein, protein levels required for normal function, or more stable epistatic protein complexes determine the penetrance of cellular fitness phenotypes throughout the course of a CRISPR experiment.

Our analysis suggests a link between the strength of ETC-related gene dependency and the screen sampling time. In the context of an effort like the DepMap, which is focused on screening large collections of diverse cell lines, there may be a complex interplay between the screen sampling time, the doubling rate of the cell line being screened, and global protein stability in each cell line (Figure 3d). While there is likely true variation in the extent of genetic dependency on mitochondria function across different cell types and genetic backgrounds, we speculate that a substantial portion of the quantitative differences observed in the strength of the ETC-related phenotypes in the DepMap may instead reflect differences in the effective sampling time, cell line doubling rate, and protein stability across these cell lines.

More generally, our study highlights the utility of FLEX, which enables objective benchmarking of functional relationships and informative summaries of the underlying functional diversity. Such methods and objective applications of them to existing data and processing methods are critical to our effective interpretation of large-scale CRISPR screens.

## Code availability

The FLEX R package can be obtained from https://github.com/csbio/FLEX_R.

## Author contribution

M.B., M.R. and C.L.M. conceived the study. M.R. and M.B. wrote the software and performed the analysis. M.R., M.B., M.C., H.N.W., K.R.B., C.B., J.M. and C.L.M. interpreted the results. A.Y.T, K.C. and M.A. performed the experiments. M.B., M.R. and C.L.M. drafted the paper with input and revisions from M.C., M.A., H.N.W., K.R.B., B.J.A., C.B., and J.M. B.J.A., C.B., J.M. and C.L.M acquired funding.

## Competing interest

J.M. is a shareholder in Northern Biologics and Pionyr Immunotherapeutics, and is an advisor and shareholder of Century Therapeutics and Aelian Biotechnology. The authors declare no conflict of interest.

## Methods

FLEX is designed to perform a systematic functional evaluation of genome-scale perturbation data. It has three different components: generation of reference standards, gene-level (global) evaluation, and module-level (local) evaluation. To use FLEX, the user must provide an input dataset and select a reference standard to evaluate against. FLEX enables both global and local functional evaluations and supports a number of visualization options (Figure S1).

### Generation of reference standards

To systematically evaluate functional relationships between gene pairs, FLEX uses various public reference datasets. These datasets include genes grouped into different modules (a set of related genes); for example, in the CORUM reference dataset, a module refers to a protein complex. Relationships between all possible gene pairs from all modules are utilized to form a co-annotation (co-membership) based binary reference standard. In such a reference standard, gene pairs co-annotated to the same module (within-module pairs) are labeled positives (1) and gene pairs from two different modules (between-module pairs) are labeled negatives (0). For all the positive pairs, the source(s) of their co-annotation (module IDs) are stored.

A single reference standard provides an exclusive view of biological complexity. To ensure functional evaluation from multiple perspectives, FLEX supports four different reference standards. For protein complexes, FLEX uses CORUM v3.0 ^8^ as the reference standard. For pathways, it uses MSigDB ^9^ that collates several pathway datasets, and for GO ^10^, it uses biological processes (BP). For reference standards based on complexes and pathways, a positive example is defined as a gene pair annotated to the same complex (or pathway). A gene pair forms a negative example when the genes come from two different complexes (or pathways). In contrast, for the GO BP dataset, FLEX first applies a filter based on the number of genes annotated to a GO term (term size). Biological processes that are too specific (term size < 10) or too general (term size >= 300) are excluded. Then using the gene annotations in the filtered GO BP candidates, FLEX applies a similar approach as outlined for complexes and pathways to define the GO BP-specific reference standard.

While all of these aforementioned reference standards are manually curated and thus high-quality, they lack in terms of their genome-wide coverage. For a broader standard, FLEX includes an integrated, data-driven reference functional network named GIANT ^11^, which reports inferred functional relationships from many different genomic or proteomic data sources. A node represents a gene in this network, and an edge represents an inferred functional relationship between two genes. While CORUM, Pathway, and GO BP provide annotations for 3662, 8904, and 13637 genes respectively, GIANT covers ~25K genes. To transform the GIANT network to a reference standard, the gene-gene relationships (edges) in the network are first ranked by the relationship strength (edge weights), in descending order. Next, the top one million gene-gene relationships are labelled as positives (the rest are negatives), resulting in a density of ~2.6% for the positive standard. The density of positives for the CORUM, Pathway, and GO BP standards are ~0.6%, ~7%, and ~6%, respectively.

Even though FLEX provides four different reference standards by default, users can easily define additional reference standards as desired. The only requirement is that the new reference standard provides associations between a subset of gene pairs. Therefore, any dataset with modules or any network quantifying gene-gene relationships is appropriate as input to FLEX to generate reference standards.

### Gene-level evaluation

FLEX performs gene-level (global) evaluations using genome-wide quantitative perturbation effects. These effects can either be dependency scores ^1^ where each gene in the library is systematically knocked out across a panel of cell lines, or genetic interaction (GI) scores ^17^, where a single gene is first knocked out and a panel of gene knockouts are introduced by library screening in the mutant background. Then, depending on user input, FLEX either calculates a pairwise profile similarity score for each gene pair (using gene profiles across screens/experiments) or uses the direct measurements between gene pairs. As FLEX evaluates gene pairs, direct measurements are only relevant when the screens also represent genes (a second knockout for GI, for example). Profile similarity scores between gene pairs are meaningful in either case (GI or dependency). FLEX uses Pearson correlation coefficient (PCC) values as measures of profile similarity.

Once a pairwise measurement for pairs of genes and their corresponding co-annotations from a reference standard are available, FLEX calculates how well the measurements agree with the co-annotations. A traditional way to capture this agreement is to use a Receiver Operating Characteristic (ROC) curve. However, as all of our reference standards are highly imbalanced (i.e. positive to negative ratio is small), a more appropriate metric to use is precision-recall ^12 18^. FLEX uses a precision-recall (PR) curve to summarize the gene-level (global) functional performance, although it plots the number of TPs (equivalent to recall on a log scale) on the x-axis instead of recall.

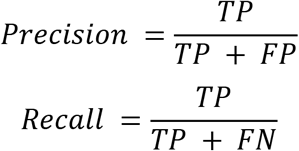

### Contribution diversity

As biological annotation standards are modularized, FLEX computes a contribution diversity matrix to understand how different modules contribute to the overall performance. Due to inherent redundancies among modules, a true positive in the standard is sometimes associated with multiple modules. To account for redundancies at the module level, FLEX estimates a subset of modules that explains all of the TPs for a set of precision levels from the gene-level PR curve. Calculating such a subset, optimally, is an NP-hard problem ^19^ and hence, FLEX uses a greedy approximation algorithm (shown below). To illustrate the algorithm (and all methods from here on), we will use CORUM as the reference standard and CORUM complexes as modules.

**Algorithm**: calculate *contribution diversity*

**Input**: a set of precision thresholds, **P**, profile similarity PCCs, CORUM standard

**Output**: a 2D matrix, **C** (row: CORUM complexes, col: precision thresholds, entry: number of TP pairs uniquely contributed by the complex at that precision threshold)

~~~
Calculate sets of TPs (also the associated complexes) for all precision thresholds, P
**for** i = 1: |P| (length of P)
        Q <- set of TPs at P[i]
        **while** Q is not empty
              Compute the number of TPs associated with individual complexes
              Rank the complexes by the number of TPs contributed (descending).
              S <- TPs associated with the highest ranked complex, j
              C[j,i] = |S| (number of contributions for complex j)
              Q <- SetDifference(Q,S) (remove S from Q)
        **end while
end for**
~~~

The algorithm outputs a precision vs complex contribution matrix that is next visualized using a Muller plot. This is termed the contribution diversity plot and it visualizes the diversity of complexes that constitute global performance. When applied to the CORUM reference standard, the contribution diversity analysis reduces the number of effective complexes to 1697 (out of 2916 total complexes), highlighting the minimal set of required complexes to explain the functional performance.

### Module-level performance

A module-level performance evaluation encapsulates the local performances of individual modules (e.g. complexes). For each complex in the CORUM reference standard, FLEX generates a per-complex subset of the reference standard that includes gene pairs between all of the genes from the complex (within-complex pairs) and pairs between genes from the complex and genes from the rest of the complexes (between-complex pairs). For each complex, within-complex pairs constitute the positive standard whereas between-complex pairs comprise the negative standard. Using these per-complex standards, FLEX calculates an area under the PR curve (AUPRC) for all individual complexes. This is visualized using a scatter plot with the AUPRC values on the x-axis and the size of the complexes on the y-axis.

### Module-level summary

FLEX generates a module-level summary plot to account for the disproportionate nature of modules in genome-wide datasets. It first calculates a module-level precision-recall (mPR) metric by computing the contribution diversity matrix (module by precision) and then counting, for each precision, the number of modules that are represented.

FLEX then visualizes this using a module-level summary plot that outputs the number of represented modules along the x-axis and precision values along the y-axis. To qualify, a complex must contribute at least one TP pair towards the performance and 10% of all of the possible within complex (TP) pairs must be present at that precision.

### Pooled CRISPR HAP1 dropout screens

Pooled CRISPR dropout screens, including CRISPR library virus production and virus titer determination, were performed as described recently ^17 20^w. In brief, human HAP1 cells were obtained from Horizon Discovery (wt: clone C631, sex: male with lost Y chromosome, RRID: CVCL_Y019) and maintained in DMEM, low glucose (10mM), 1mM glutamine, 10% FBS.

CRISPR library virus production was performed in HEK293T cells. Therefore, 10 million cells were seeded per 15 cm plate in DMEM medium containing high glucose, pyruvate and 10% FBS. Twenty-four hours after seeding, the cells were transfected with a mix of 8 μg lentiviral lentiCRISPRv2 vector containing the TKOv3 gRNA library ^21^ (Addgene #90294), 4.8 μg packaging vector psPAX2, 3.2 μg envelope vector pMD2.G, 48 μl X-tremeGene 9 transfection reagent (Roche) in 1.4 ml Opti-MEM media (Life Technologies) for a total volume of 800 μl. Virus-containing media was harvested 48 hours post transfection.

For pooled CRISPR dropout screens, 3 million HAP1 cells were seeded in 15 cm plates in 20 ml of specified media. A total of 50-90 million cells were transduced with the lentiviral TKOv3 library at a MOI~0.3, so that each gRNA is represented in about 200-300 cells. 24 h post infection, transduced cells were selected in 1 μg/ml puromycin for 48 hours. Cells were then harvested and pooled, and 30 million cells were collected for subsequent genomic DNA extraction and determination of the library representation at day 0 (i.e. T0 reference). The pooled cells were seeded into three technical replicate plates, each containing 15 million cells (>200-fold library coverage) and passaged every three to four days and at >200-fold library coverage until T18. Cell pellets from each replicate were collected at each timepoint of cell passage.

Genomic DNA was extracted using the Wizard Genomic DNA Purification Kit (Promega). Sequencing libraries were prepared from 50 μg of the extracted genomic DNA in two PCR steps, the first to enrich guide-RNA regions from the genome, and the second to amplify guide-RNA and attach Illumina TruSeq adapters with i5 and i7 indices. Barcoded libraries were gel purified and final concentrations were estimated by quantitative RT-PCR. Sequencing libraries were sequenced on an Illumina HiSeq2500 instrument using single-read sequencing. The T0 and T18 time point samples were sequenced at 400- and 200-fold library coverage, respectively.

### Mapping of reads to gRNAs

FASTQ files from single-read sequencing runs were first trimmed by locating constant sequence anchors and extracting the 20 bp gRNA sequence preceding the anchor sequence. Trimmed reads were aligned to the TKOv3 library reference using Bowtie (v0.12.8) allowing up to 2 mismatches and 1 exact alignment (specific parameters: -v2 -m1 -p4 --sam-nohead). Successfully aligned reads were counted, and merged along with annotations into a matrix.

### LFC precision-recall analysis

To control quality of genome-wide CRISPR/Cas9 screens in HAP1 cells, gene-level fitness effects were estimated by first computing a log2 fold-change (LFC) quantifying the dropout of a gRNA from the population between T0 (after puromycin selection) and a given mid or end point (T3 - T19). The LFC values of the four gRNAs targeting a given gene were mean summarizing. Gold-standard essential (reference) and non-essential (background) gene sets were taken from Hart et al., 2015 ^22^ and Hart et al., 2017 ^21^. For the identification of reference (essential) genes using LFC values of a given screen was assessed by computing precision over true positive statistics.

### Calculation of ranks for HAP1 screen time points

Using the common essential and non-essential gene sets that were used for scaling the DepMap CERES scores, we scaled the gene dropout effects in HAP1 cells. At each time point between T3 and T19 LFC values, which represent the difference between a given time point and T0, were scaled to ensure a median score of −1.0 for the essential genes and 0 for the non-essential genes. We then merged each of the 31 HAP1 screen time points with the 563 Broad DepMap screens and calculated median scores for a subset of genes (genes from ETC-related complexes, spliceosome, 26S proteasome, cytoplasmic ribosome). Finally, we ranked all HAP1 screens (time points between T3 and T19) separately by the calculated median score. The correlation between the resulting ranks for all 27 screens sampled between T6 and T19, and the time points at which a given screen had been sampled was computed using a Wilcoxon rank-sum test. Note that due to strongly varying drop out patterns of core essential genes as well as strongly variable LFC values for non-essential genes, LFC scaling generated several extreme values. Therefore, to computing more conservative correlations coefficients between sampling time and ranking, the four T3 screens were excluded.

### Estimating the gene and protein complex dropout speed

For each of the 71k gRNAs in the TKOv3 library, 31 LFC measurements were taken between T3 and T19 in wildtype HAP1 cells. A loess model was fit through the 31 measurements and T0, estimating the interpolated LFC at 0.2 days resolution. All possible differential (d)LFC values were computed by contrasting interpolating LFC values 3 days apart:

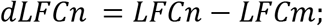

where n is between T3 and T19, m is between T0 and T16, and n - m is equal to three days. Furthermore, LFC is the interpolated LFC value for a given gRNA. Since absolute dLFC values depend on the maximal dropout (Figure S8d, e), the time point of the maximal dropout was estimated by maximizing the separation of non-essential and core essential gene LFC values. The gRNA-level dependency of dLFC values on the LFC at this maximum dropout point was removed by computing the residuals from a Loess fit (Figure S8d, e). For each gene, gRNA residuals are mean summarized at each point between T3 and T19 to define the dropout speed. For each of the CORUM complexes, the respective gene-level dropout speed was median summarized.

---

**Figure S1:**
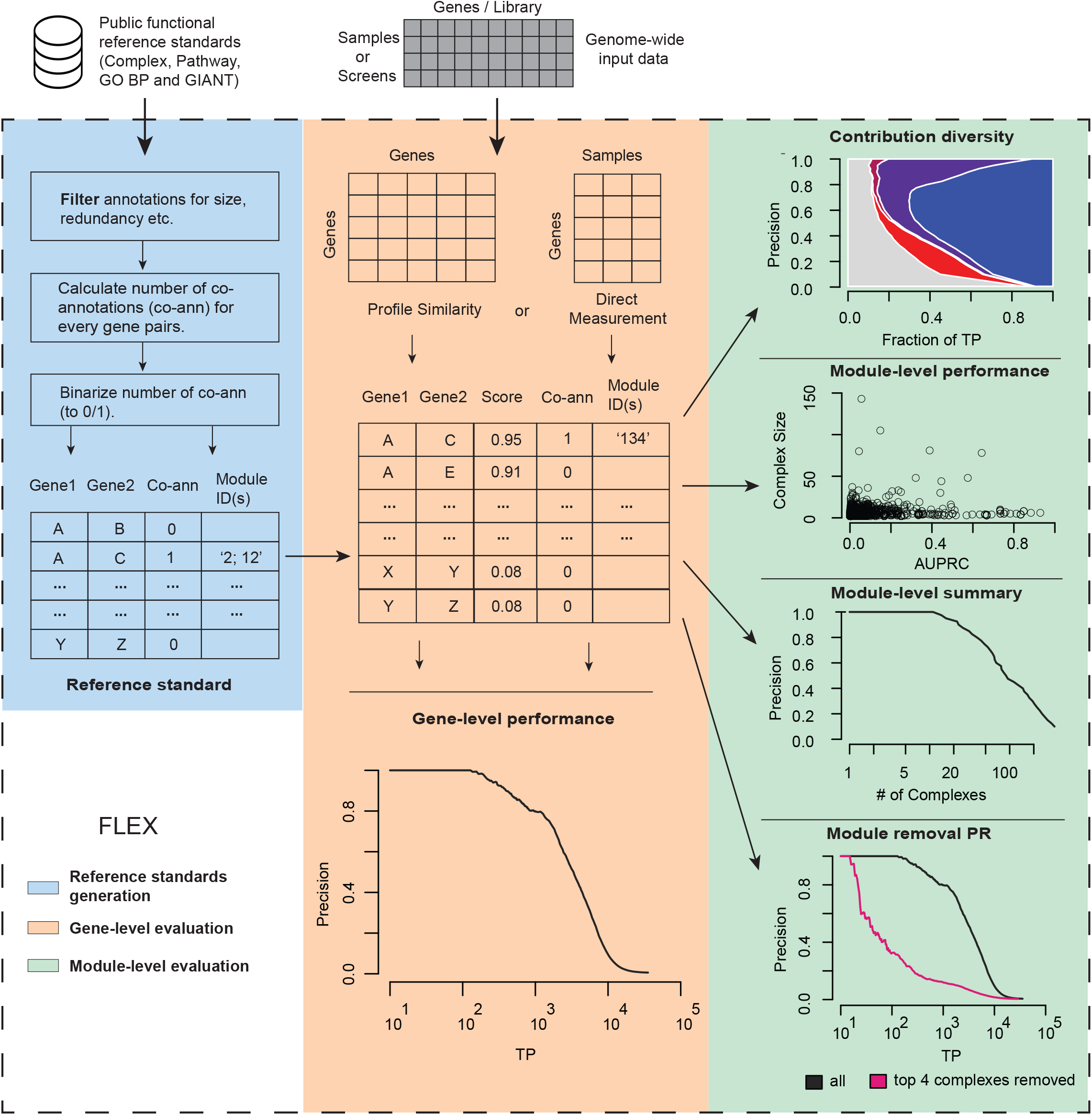
Overview of FLEX. FLEX investigates and summarizes the functional information in an experiment. **Reference standards generation (light blue)**. FLEX takes as input public functional reference standards and derives a gene-pair binary reference standards for downstream analysis. **Gene-level evaluation (light orange)**. FLEX generates a PR curve summarizing the gene-level (global) performance of the experimental data on CORUM co-memberships. The experimental data can either be dependency scores or genetic interactions. **Module-level evaluation (light green)**. FLEX interprets the module-level (local) functional signal of CORUM complexes. *The contribution diversity* plot summarizes the diversity of protein complexes that contributes to the global PR performance. The top four complexes are highlighted. *The module-level performance* plot visualizes the performance of individual protein complexes. *The module-level summary* plot lists the number of complexes captured at different precision levels. *The module removal PR* plot explains functional bias in global PR performance. The top four complexes are removed from the data and gene-level PR-performance is re-evaluated (pink line).

**Figure S2:**
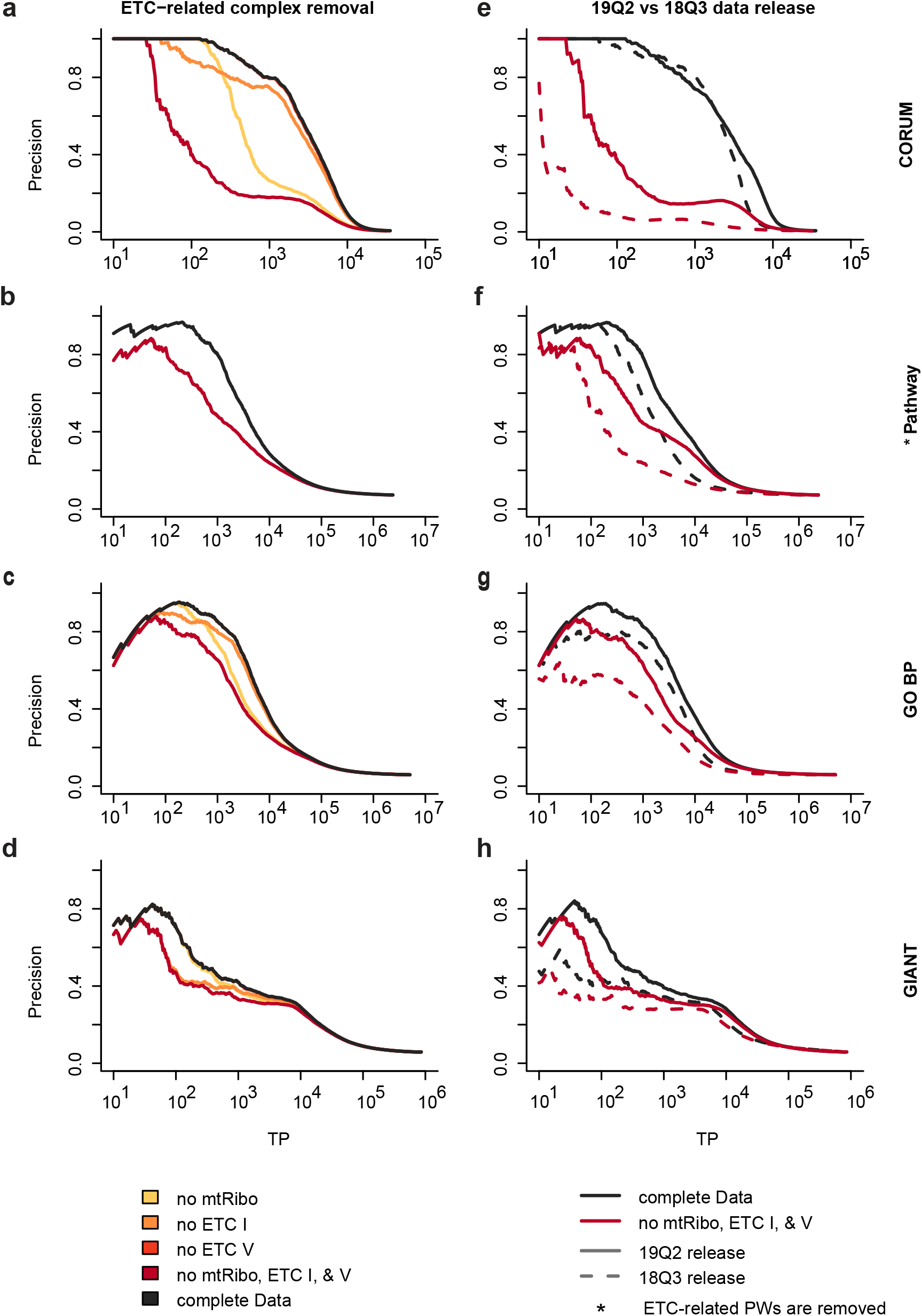
Mitochondrial complex removal effect on performance of different DepMap CERES score versions across reference standards. Precision recall curves summarize the performance of DepMap co-essentiality profiles to capture functional relationships reported in different standards. **a-d**, Performance decrease of the DepMap 19Q2 release CERES scores after removal of ETC-related complexes. The ETCI (orange), ETCV (light red) and 55S mitochondrial ribosome (yellow) are removed individually and together (dark red) for CORUM, GO BP, and GIANT standards. For Pathway, ETC-related pathways (dark red) are removed (as the Pathway standard does not contain members of the 55S mitochondrial ribosome). Shown is the performance on the standards CORUM 3.0 complexes (a), Pathway (b), Gene Ontology (GO) Biological processes (BP) (c), and the GIANT functional network (d). **e-h**, The performance of DepMap CERES score 18Q3 and 19Q2 releases are compared on the four above-mentioned standards. ETC-related complex removal affects the 18Q3 release performance more strongly than the 19Q2 release performance. The same sets of cell lines from 19Q2 and 18Q3 are used for this comparison.

**Figure S3:**
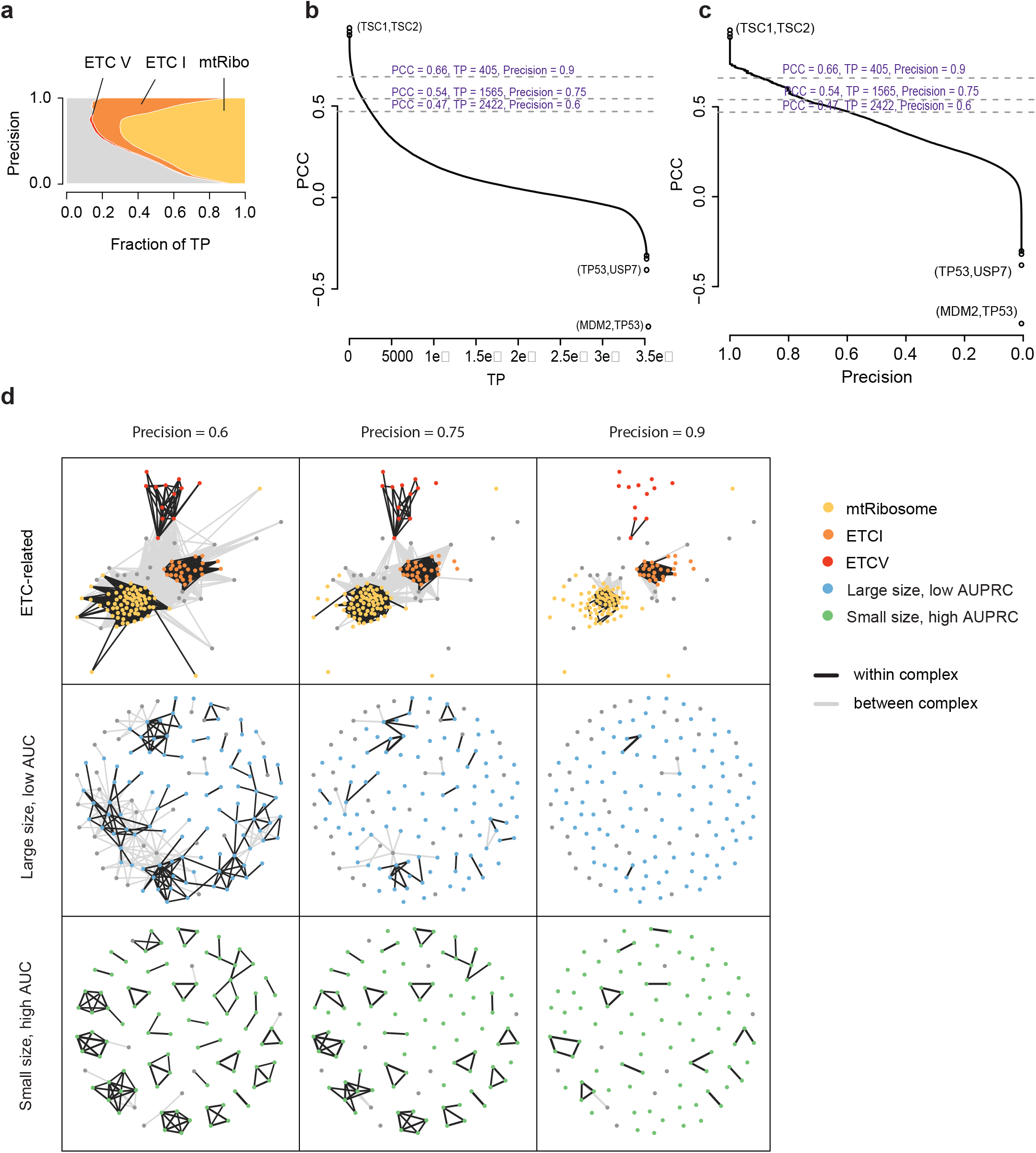
Protein complex PR contribution. **a**, Contribution diversity of CERES score PR performance using the CORUM complex standard with a focus on ETC-related complexes. Shown are the fraction of TP pairs for CORUM complexes at different precision thresholds. **b**, PCC vs TP plot for gene pairs co-annotated to the CORUM complexes. Corresponding precision values for a subset of PCC and TP values are shown. **c**, Similar to **b**, but Precision values are plotted on the x-axis. **d**, Co-essentiality networks using different PCC values as cutoffs (selected at different precision values of 90%, 75% and 60%) and are grouped by ETC-related genes, large size, low AUC genes, and small size, high AUC genes. Gene pairs from within the same complex (TP or co-annotated) are connected through black edges and between complex (FP) gene pairs are connected by light gray edges. Large size, low AUC genes are from complexes with more than 30 members and AUC smaller than 0.4. Small size, high AUC genes are from complexes with less than 30 members and AUC larger than 0.4. ETC-related genes, large size, low AUC genes, and small size, high AUC genes and are color-coded.

**Figure S4:**
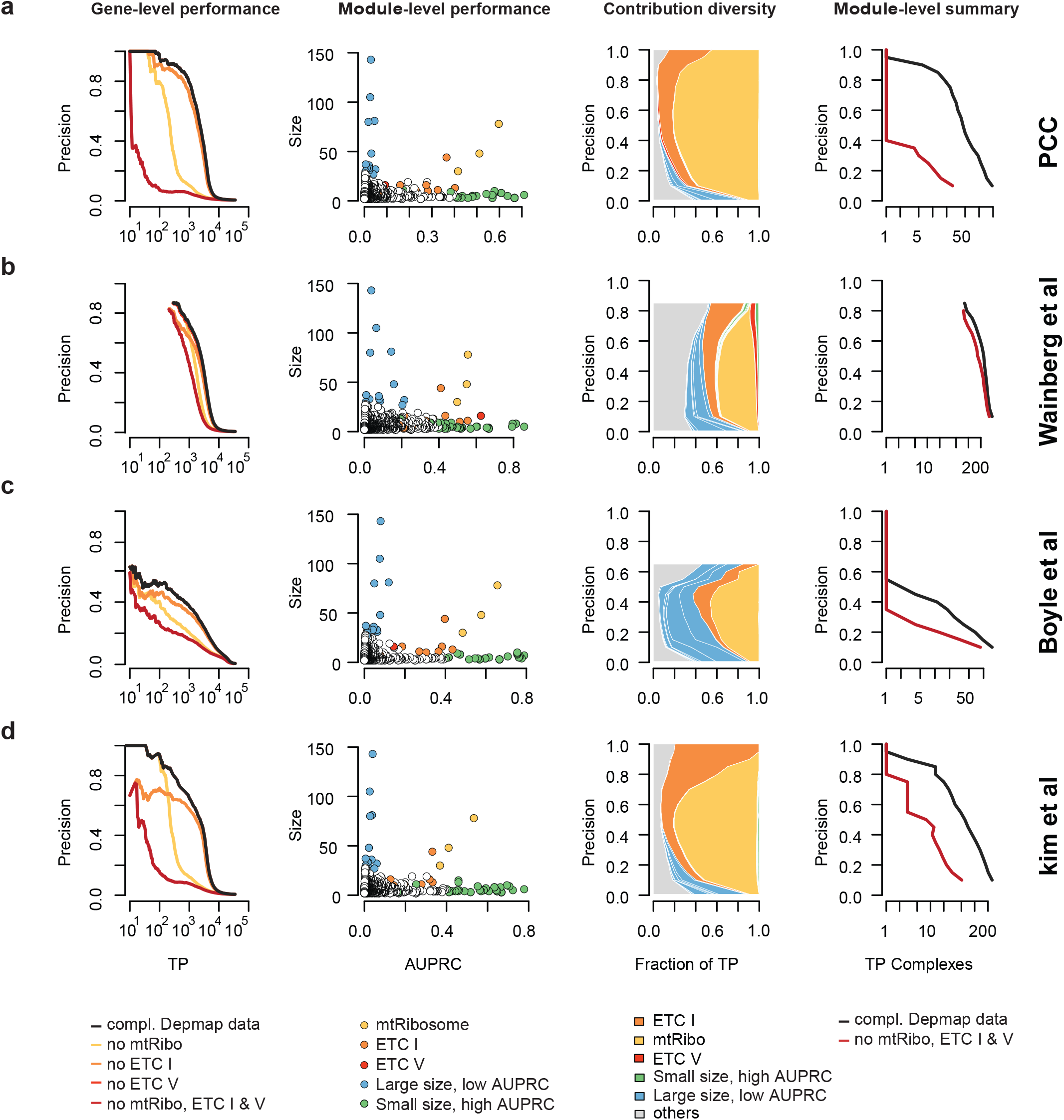
FLEX exploration of alternative DepMap-to-network computational approaches. Explored are different alternative methods to infer functional relationships from DepMap 18Q3 release CERES score co-essentiality profiles and CORUM 3.0 complex relations are used as standard. **a,** Pearson correlation coefficient (PCC) to infer relationships. **b,** Generalized least squares (GLS) based approach by Wainberg and colleagues ^7^. This approach bases gene pair similarity scores on FDR corrected p-values (1 - fdr) resulting in a ‘late start’ of the PR curve (many values at top are the same, 1.0). **c,** CERES score matrix multiplication normalization using olfactory genes to estimate noise in the data by Boyle and colleagues ^4^. **d,** PCC-based similarity approach preceded by gene and screen filtering by Kim and colleagues ^6^.

**Figure S5:**
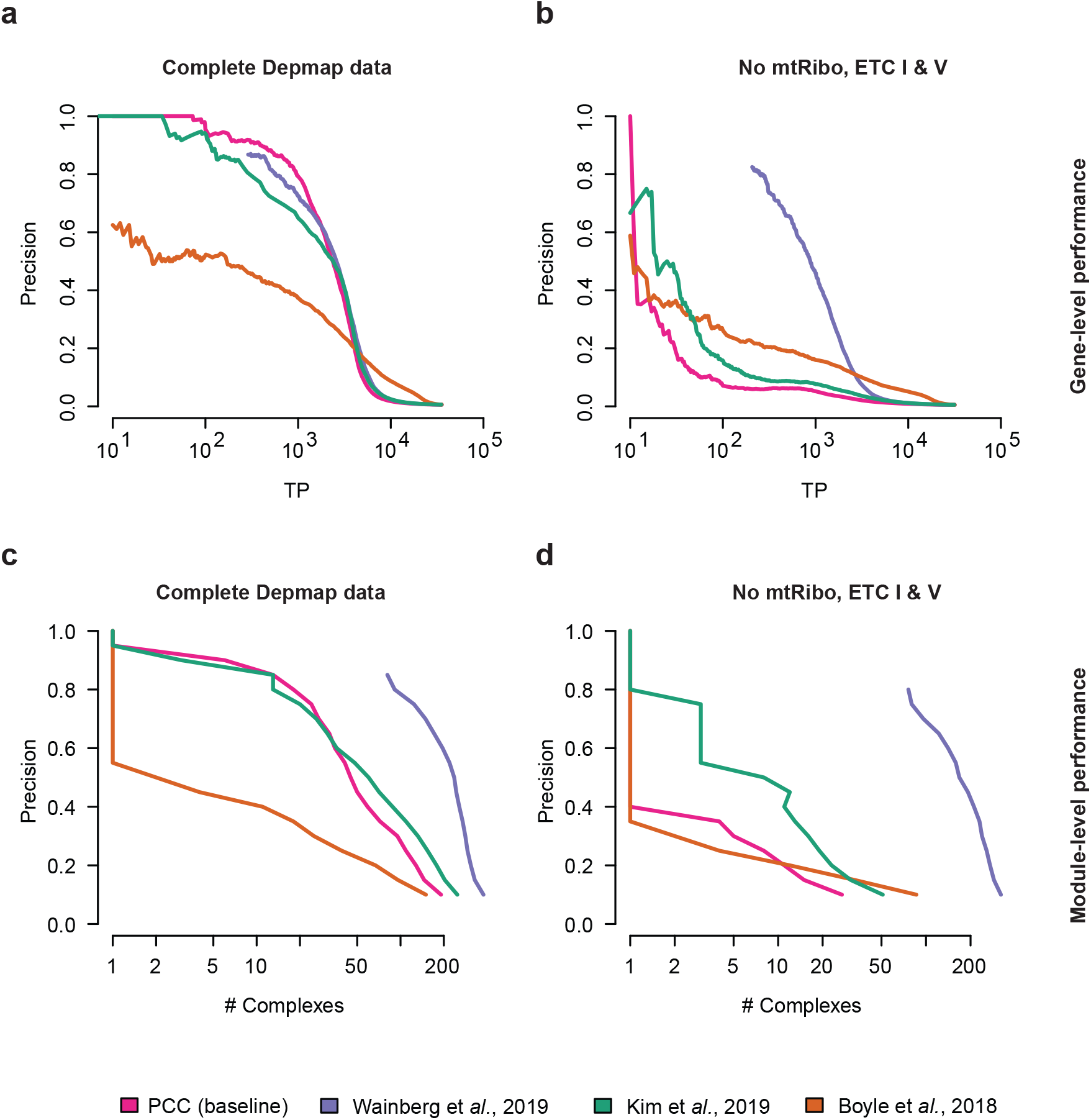
Direct comparison of alternative DepMap-to-network computational approaches. Compared are different alternative methods to infer functional relationships from DepMap 18Q3 release CERES score co-essentiality profiles and CORUM 3.0 complex relations are used as standard. The methods are PCC of co-essentiality profiles as baseline data treatment (red) and approaches published in Wainberg and colleagues ^7^ (violet), Boyle and colleagues ^4^ (orange) and Kim and colleagues ^6^ (green). **a,** Gene-level performance to capture co-complex membership on the full data. **b,** Gene-level performance to capture co-complex membership when ETC-related complexes are removed from data and standard. **c,** Module-level performance to capture number of complexes with TP co-complex membership on the full data. **d,** Module-level performance to capture number of complexes with TP co-complex membership when ETC-related complexes are removed from data and standard.

**Figure S6:**
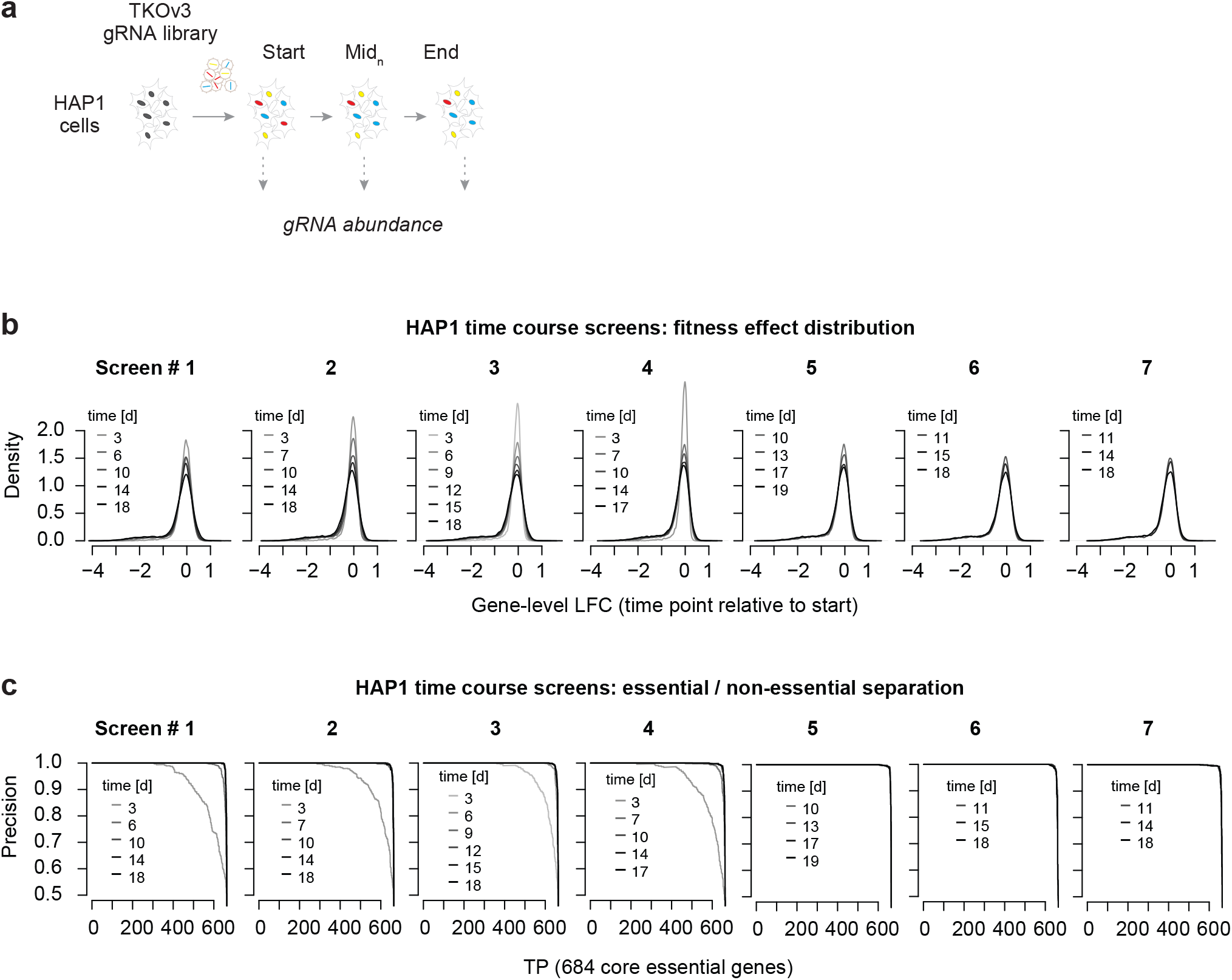
HAP1 genome-wide CRISPR screen time course data quality. **a**, Schematic illustration of the experimental workflow for time-resolved genome-wide CRISPR/Cas9 screens in HAP1 cells. **b**, Fitness effect distribution of 17,804 genes targeted with the TKOv3 gRNA library at different time points in the 7 independent screens. The fitness effect is measured by computing a log2 fold-change (LFC) of gRNA abundance at a given time point compared to the starting population (T0; after puromycin selection). **c**, Screen quality control for all time points of the 7 independent screens. This was done by testing the capacity of LFC values to separate 684 core essential and 927 non-essential genes (see Methods).

**Figure S7:**
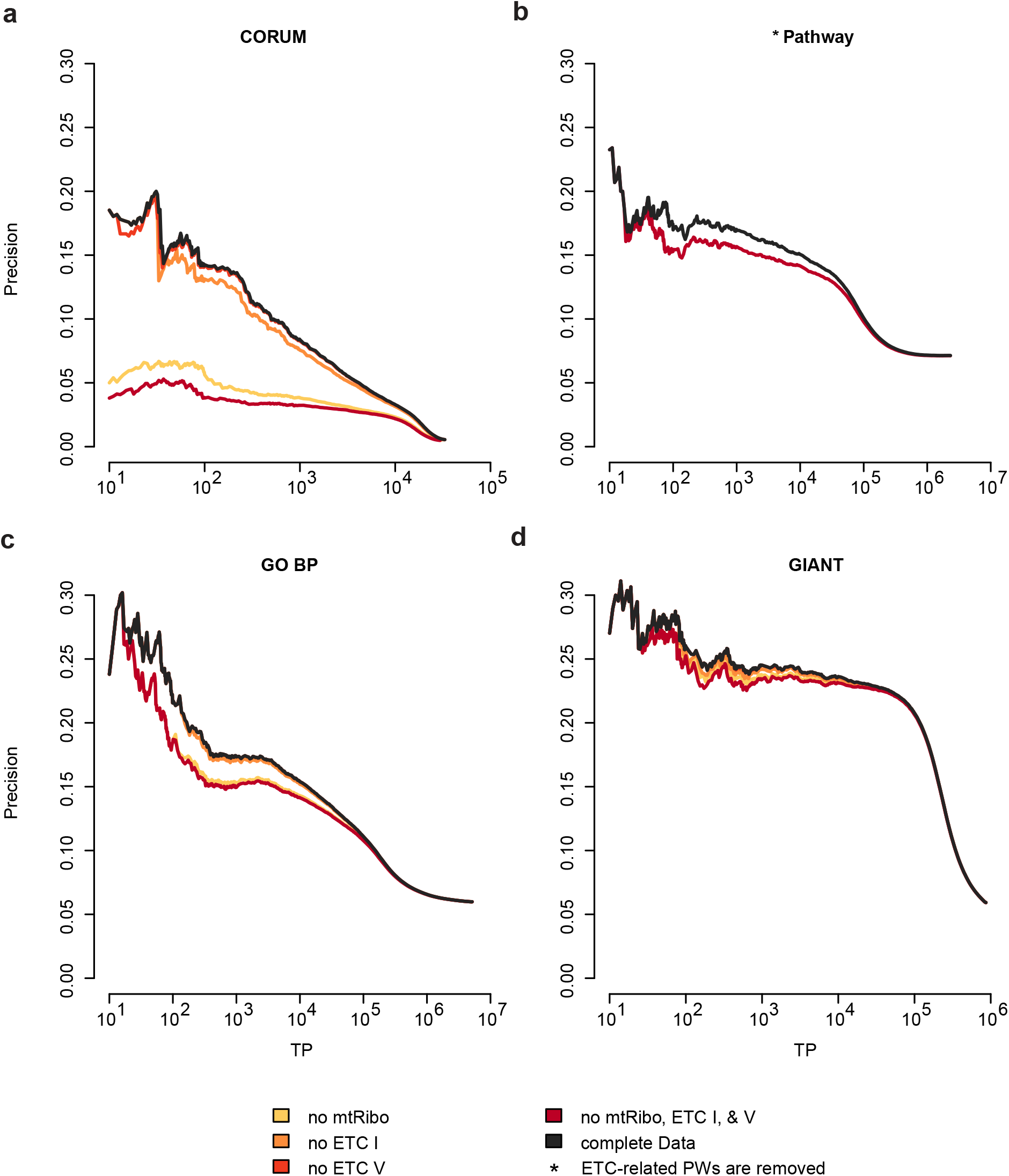
Time-resolved CRISPR screens in a single (HAP1) cell line contain a dominant ETC-related functional signature. PR performance of time-course fitness effect profiles in HAP1 cells to capture relationships from CORUM complexes (a), Pathways (b), GO BP (c) and GIANT (d). Pearson correlation coefficients (PCC) are computed between interpolated LFC profiles along the HAP1 time-course. Black line shows complete data, orange, light red and yellow lines show performance after ETC I, V and 55S mitochondrial ribosome removal, respectively. The dark red line shows performance upon removal of all three complexes from the data and standard for CORUM, GO BP, and GIANT. For Pathway, the dark red line shows performance upon removal of ETC-related pathways (the Pathway standard does not contain members of the 55S mitochondrial ribosome).

**Figure S8:**
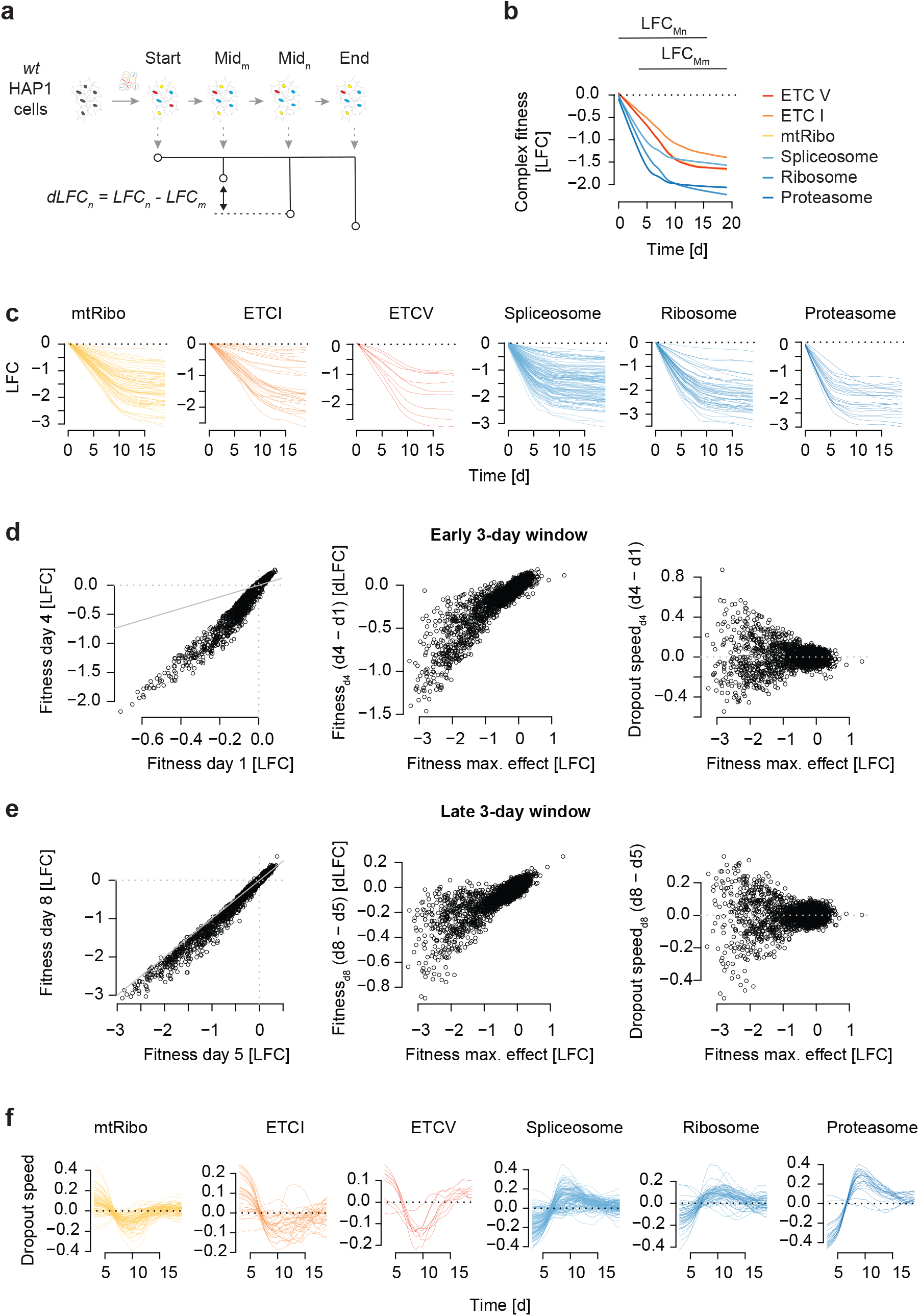
Temporal fitness phenotype estimation on HAP1 genome-wide CRISPR screening data. **a**, Schematic illustration of data processing to estimate dropout rate across time. The contrast of a pair of sliding intermediate time points (Mn, Mm) is used to estimate drop-out rates. **b**, Fitness effects of selected complexes along the time of a screen. Shown is the median gene dropout LFC across a given complex. LFC values for each gRNA are first interpolated from 31 measurements derived from 7 independent screens between 3 and 19 days past puromycin selection (screen start). Next, gRNA-level interpolated LFC values are mean-summarized to the gene-level and finally median-summarized to the complex-level as defined by CORUM 3.0. The range of the sliding intermediate time points is indicated on top. **c**, Fitness effects of the genes in a selected complex along the time of a screen. **d**, **e**, Dropout rate estimation in an early (d) and mid/late (e) window along time. Window size is 3 days (day 4 - day 1 or day 8 - day 5). Gene-level fitness effects (LFC) comparison between sets of time points (left). The differential LFC values (y-axis) depend on LFC values measured at the time point with the strongest phenotypic effect (T18; see methods) (middle). This dependency is normalized for to estimate a dropout rate, which is independent of a gene's fitness phenotypic strength (right). Normalization is done by taking the residual from the loess fit. **f,** Adjusted differential fitness effect (negative dropout speed) of genes for each of the selected complexes shown in Figure 2h.

**Figure S9:**
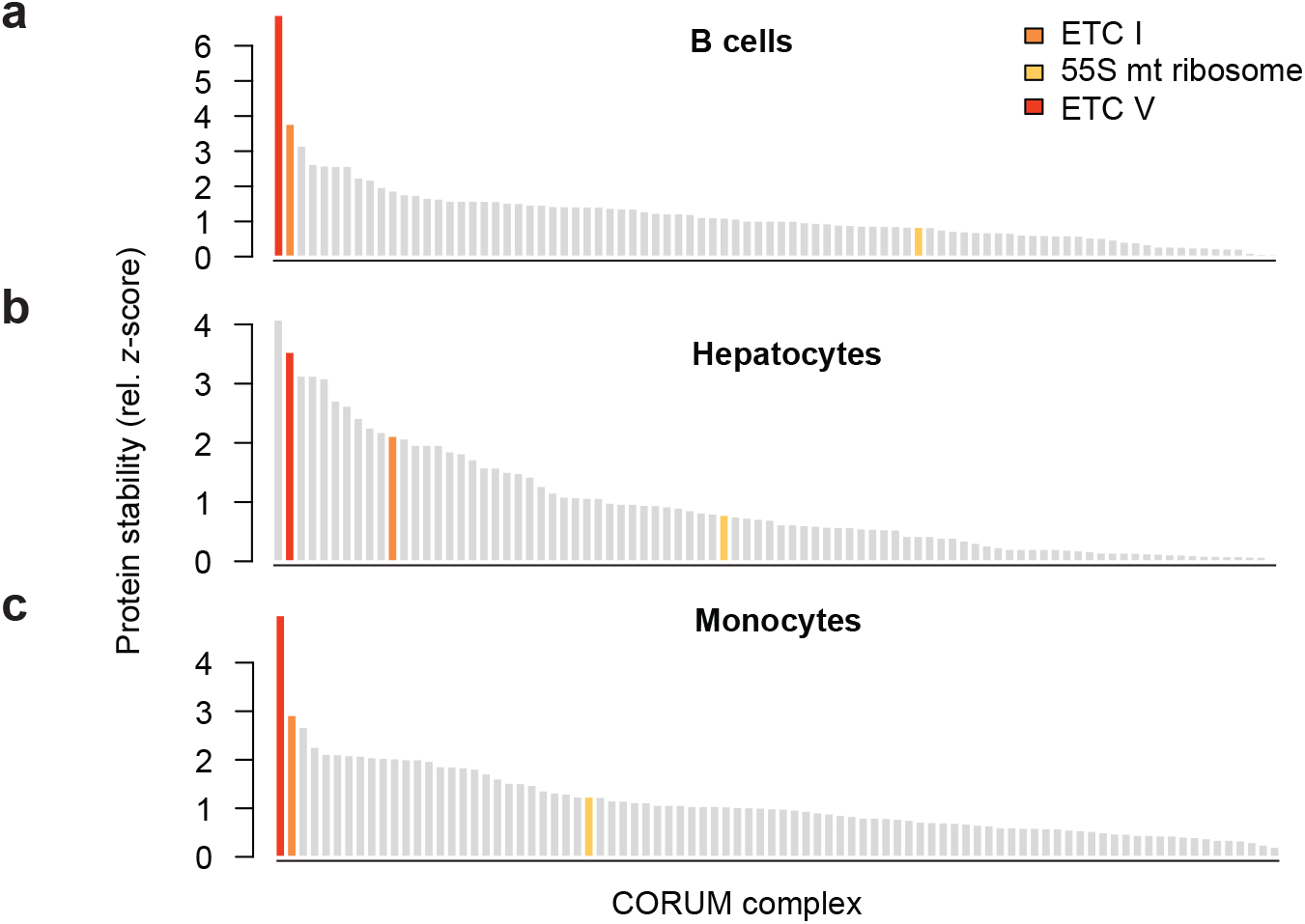
Stability of CORUM protein complexes across three cell lines. Shown is the data for B cells (a), hepatocytes (b) and monocytes (c). Protein half-life data was taken from Mathieson et al., 2018 14. The data was summarized on CORUM 3.0 complex level. Half-life data was z-transformed, and the minimum z-score set to 0 to emphasize large z-scores. Complexes for which at least 5 members contributed data across the three cell lines are shown.

**Figure S10:**
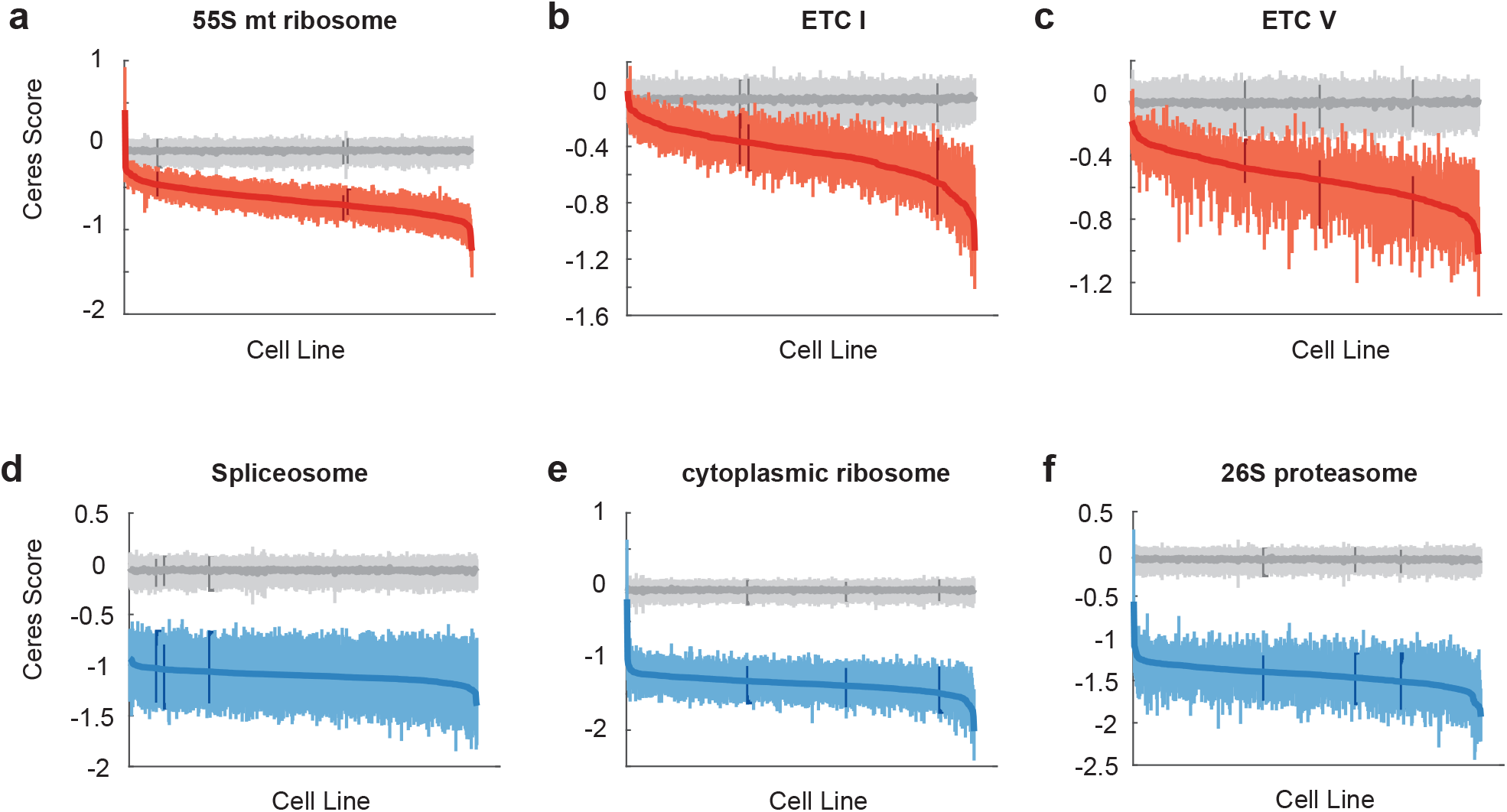
DepMap CRISPR screens (release 19Q2) sorted by complex CERES score. DepMap genome-wide CRISPR screens ranked by the median CERES score across the 55S mitochondrial ribosome (a), the ETC I (b) and V (c) as well as across selected essential complexes including spliceosome (d), cytoplasmic ribosome (e) and 26S proteasome (f). The middle blue/red line indicates the median, the vertical lines the 25% and 75% quantiles of a given screen. Grey lines represent the same metrics for all genes in the genome.

